# Dysregulation of miRNAs Drives Premature GABAergic Maturation and Early Neurodevelopmental Defects in Schizophrenia

**DOI:** 10.64898/2026.05.25.727574

**Authors:** Fadumo A. Mohamed, Rolf Søkilde, Elif Bayram, Arkadiusz Nawrocki, Pia Jensen, Marion Kadlecova, Methi Wathikthinnakon, Susanna Cirera, Trine Møller, Charlotte Brasch-Andersen, Michael Benros, Martin R. Larsen, Boye S. Nielsen, Kristine Freude

**Author notes:** Contributed equally (Fadumo A. Mohamed, Rolf Søkilde). Corresponding author: Kristine Freude.

## Abstract

**Background:** Schizophrenia (SCZ) is a severe neurodevelopmental disorder with numerous genetic risk loci. However, little is known about the molecular alterations that occur during brain development in SCZ, particularly regarding the role of microRNA (miRNA) mediated regulatory mechanisms. This gap in knowledge is largely due to the limited availability of developing human brain tissue. Patient-derived brain organoids offer a promising alternative model. Here we use 3D dorsal forebrain organoids (DFOs) to investigate miRNA dysregulation in SCZ.

**Methods:** DFOs were generated from human induced pluripotent stem cells (hiPSCs) derived from six SCZ patients and five matched controls and cultured for 120 days. Multi-omics analyses, immunohistochemistry, and in situ hybridization were employed to characterize molecular and spatial features.

**Results:** DFOs recapitulated key molecular hallmarks of human cortical development. Nineteen miRNAs were differentially expressed in SCZ: nine associated with neural progenitor proliferation were downregulated and ten linked to neuronal differentiation and synaptic maturation were upregulated, reflecting a compressed developmental timeline. Among 77 dysregulated mRNAs, 55 were predicted miRNA targets. SCZ DFOs exhibited significant upregulation of GABAergic pathway genes accompanied by altered expression of their regulatory miRNAs, indicating premature GABAergic lineage specification. The disrupted miRNA–mRNA network converged on glutamatergic and dopaminergic development, synaptic organization, and extracellular matrix remodeling.

**Conclusion:** Dysregulated miRNAs in SCZ DFOs disrupt neuronal differentiation, excitatory–inhibitory balance, and early circuit formation, implicating miRNA-mediated post-transcriptional regulation as a key mechanism linking molecular alterations to cellular and network-level deficits in SCZ.

## INTRODUCTION

Schizophrenia (SCZ) is a severe neurodevelopmental disorder that typically manifest from late adolescence or early adulthood [1]. Symptoms include hallucinations, delusions, social withdrawal, cognitive deficits, and mood disturbances. There is a strong genetic component to SCZ, with over 287 risk-associated loci and 106 protein-coding genes identified to date [2]. Additionally, epigenetic factors like miRNAs are implicated in the pathophysiology of this polygenic disorder [3]. These small non-coding RNAs regulate gene expression post-transcriptionally and by targeting hundreds of mRNAs leading to widespread disruption of protein interaction networks and cellular pathways [4]. Transcripts of SCZ-associated genes harbor more predicted miRNA binding sites than those of non-associated genes, implying enhanced miRNA-mediated regulation that may influence SCZ pathophysiology [5, 6]. Concordantly, multiple miRNAs, including miR-137, miR-195, miR-346, and miR-103a, are dysregulated in SCZ and relate to cognitive phenotypes and antipsychotic exposure, supporting their biomarker potential. Postmortem studies further show that ∼19% of miRNAs exhibit diagnosis-related reductions [7–9]. However, postmortem studies provide only static snapshots of late-stage disease, and miRNAs detected in blood samples may not accurately reflect those expressed in the brain, as their levels can be influenced by factors such as medication and disease duration [5]. Therefore, miRNAs identified in patient or postmortem tissues may not capture their dynamic regulation during neurodevelopment. Thus, hiPSC-derived brain organoids provide a suitable model to study developmentally regulated miRNAs, recapitulating key genetic, cellular, and molecular features of early human brain development [10].

Only few studies have examined dynamic miRNA expression during neurodevelopment in the context of schizophrenia [11, 12], and prior SCZ organoid work has generally profiled mRNA or protein at single time points without assessing miRNA dynamics or aligning all three molecular layers across development [13, 14]. To address this gap, we generated dorsal forebrain organoids (DFOs) from SCZ and matched control hiPSCs and conducted longitudinal multi-omics profiling across three maturation stages. Although SCZ and control DFOs followed broadly similar differentiation trajectories, SCZ DFOs showed consistent alterations in miRNA expression, gene regulatory programs, and protein composition that converged on neuronal maturation, synaptic development, neurotransmitter signaling, and extracellular matrix organization. To our knowledge, this represents the first comprehensive integration of miRNA, mRNA, and protein regulation during human neurodevelopment in SCZ using DFOs as a tractable model of early, cell-intrinsic disease mechanisms.

## METHODS AND MATERIALS

### See Supplemental Methods (SM) for details of all methods

#### hiPSC Generation and Characterization

Ethical approval (S-20220037) and recruitment are described in SM. After obtaining informed consent patients were included and hiPSCs were generated from the derived PBMCs from 6 SCZ and 5 age- and sex-matched healthy control (CTRL), median age 21. The WHO Schedules for Clinical Assessment in Neuropsychiatry (SCAN, version 2.1) interview was used to validate schizophrenia diagnoses given to patients by clinical doctors prior to inclusion and to rule out prior or current mental disorders in the healthy controls.

#### Generation of DFOs

DFOs were generated from all hiPSC lines using a Dorsal Forebrain Organoid kit (Cat. No. 08620, Stemcell Technologies). DFOs were differentiated until day (D)120 and analyzed at D40, D80, and D120.

#### Sample Fixation and Immunohistochemistry

Two DFOs per cell line and timepoint were fixed in either 4% paraformaldehyde or 10% neutral buffered formalin. Details on immunohistochemistry are listed in SM.

#### In Situ Hybridization

Automated in situ hybridization was performed using Locked nucleic acid (LNA) probes as previously described [15, 16] with details listed in SM.

#### Microscopy

hiPSCs and DFOs were imaged with a Zeiss LSM 700 confocal microscope, Panoramic confocal slide scanner (3DHISTECH) or Zeiss Axioscan.Z1.

#### RNA-Seq and qPCR for miRNA and mRNA Analysis

RNA from five DFOs per cell line at D40, D80, and D120 was extracted (miRNeasy Micro Kit). Library prep (Lexogen/NEBNext) and sequencing (NovaSeq X Plus, 150 bp) were performed by Omiics ApS. Small RNAs were mapped to tRNAs, miRNAs, other small RNAs, mRNA/rRNA, and the genome. Read counts (Ensembl v112) were provided by Omiics; Entrez IDs were added for analysis. qPCR validation of 19 miRNAs and 4 mRNAs was performed, using two reference genes [17, 18].

#### Proteomics

For each time point, protein was isolated from 3-5 pooled organoids. Protein concentration was measured before reduction with DTT, alkylation, and digestion. Peptides were acidified, centrifuged, loaded onto Evosep C18 tips then separated (Evosep One) before analysis by dia-PASEF (Bruker timsTOF Pro) and data processing. Differential expression analysis followed the same DESeq2 model as used for miRNA and mRNA data.

## RESULTS

### Validation of Neurodevelopment and DFO Identity

DFOs were cultured for 120 days, with samples collected on D40, D80, and D120 for multi-omics profiling to assess miRNA, mRNA, and protein expression. DFO development was validated *via* morphological assessment and marker expression analysis. At D40, DFOs expressed neural progenitor (SOX2) and neuronal markers (β-TUBULIN, MAP2) (Figure 1A). At D80, dorsal forebrain identity was confirmed by the co-expression of PAX6, FOXG1, and TBR2, along with the neuronal marker β-TUBULIN (Figure 1B). By D120, the DFOs expressed markers indicative of cortical neuron development, including deep-layer (CTIP2) and upper-layer (SATB2) markers (Figure 1C), as well as glial markers (GFAP and S100B) (Figure 1D). Simultaneously, the mature neuronal marker MAP2 was expressed, indicating ongoing developmental maturation consistent with expected cortical development.

**Figure 1.**
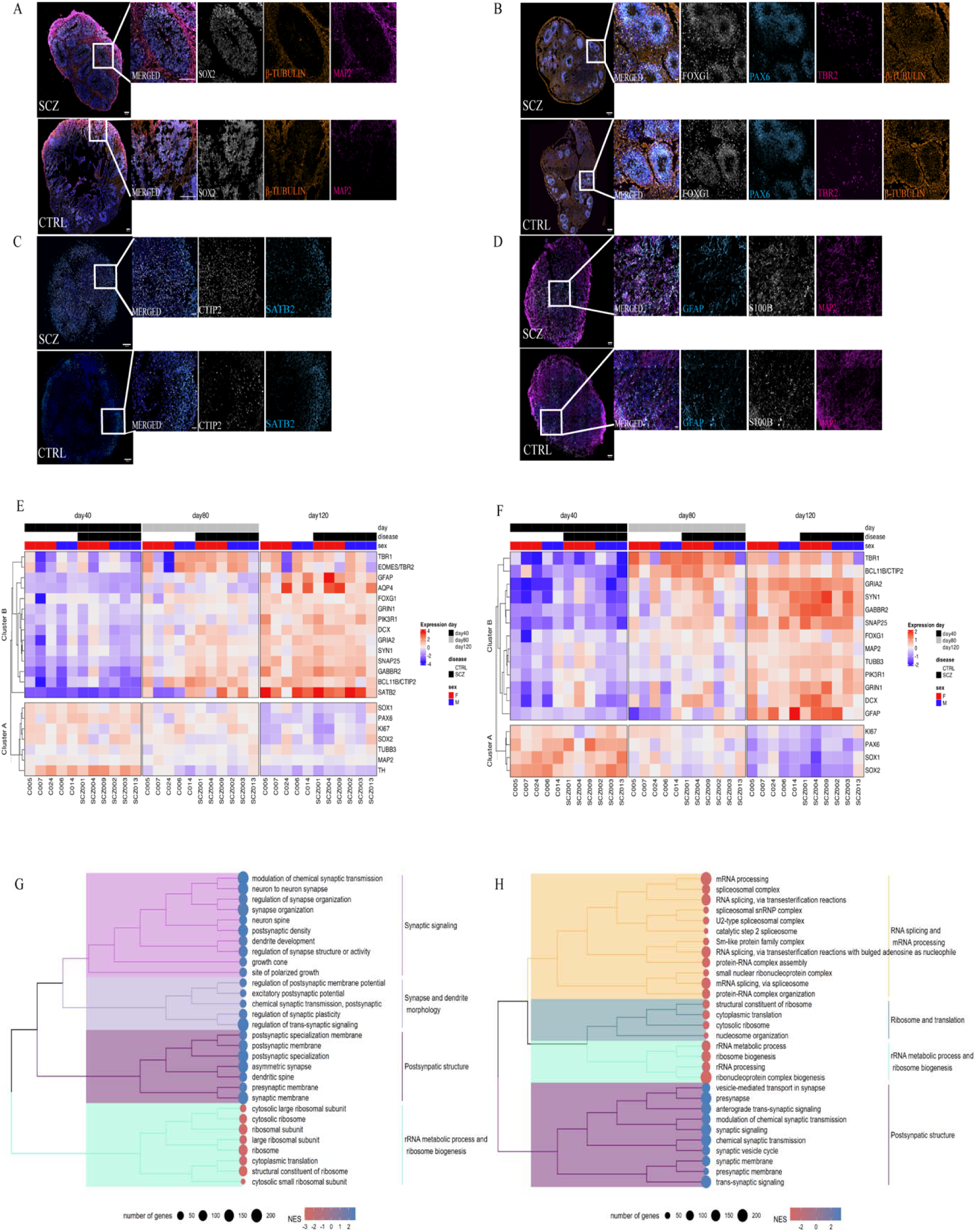
Molecular and cellular features confirm neurodevelopmental trajectory and dorsal forebrain identity of organoids: **A.** Confocal images of D40 DFOs show early neurogenesis: neural rosettes with SOX2 marks progenitors, β-tubulin marks early neurons, and MAP2 marks mature neurons. DFO: SCZ001 and C014. **B.** D80 DFOs express dorsal forebrain markers: FOXG1, PAX6, and TBR2 in distinct spatial patterns indicating active neurogenesis and DFO identity. DFO: SCZ004 and C007 **C.** Cortical layer at D120 shown by CTIP2 (deep layers) and SATB2 (upper layers); partial overlap reflects early cortical specification. **D.** At D120 DFOs express astrocytic markers GFAP and S100B alongside MAP2, indicating mature neurons and astrocytes. SCZ009 and C005 are shown. For all staining DFOs generated from the other cell lines can be found in Supplementary Figure 1-5. **E–F.** Heatmaps show consistent dorsal forebrain and cortical marker expression at mRNA (E) and protein (F) levels. Cluster A shows NPC markers; cluster B shows neuronal differentiation markers.

Gene expression profiles (Figure 1E) and protein expression heatmaps (Figure 1F) demonstrated developmental trajectories characteristic of DFOs across all cell lines. Proliferative markers such as KI67, SOX1, SOX2, and PAX6 showed high expression at D40 and decreased by D120. Conversely, markers of mature neurons (DCX, SYN1) and DFO-specific markers (FOXG1, TBR1) were upregulated at both D80 and D120, reflecting expected progression of neurodevelopment. Despite minor variability between individual DFOs and hiPSC lines, overall developmental trajectories were consistent. GO term analyses of early (D40) vs. late (D120) stages showed increased expression of genes involved in synaptic signaling and postsynaptic structure, with concurrent downregulation of rRNA metabolism and ribosome biogenesis (Figure 1G, H), indicating a shift from proliferation to synaptic maturation. These findings confirm that, despite minor variability between individual DFOs and hiPSC lines, our protocol reliably generates DFOs with comparable developmental progression in both SCZ and CTRL following overall developmental trajectories

### miRNA Expression Dynamics Reflect Neurodevelopmental Progression in DFOs

DFO maturation was accompanied by dynamic, stage-specific shifts in miRNA expression (Figure 2). Differential expression analysis of miRNAs between D40 and D120 DFOs, irrespective of SCZ or CTRL status, identified 163 downregulated miRNAs linked to pluripotency and proliferation, and 76 upregulated miRNAs involved in neural differentiation and development (Figure 2A-B). Initial upregulation of the neurodevelopmental regulator miR-9 and the subsequent decline aligns with the transition from progenitor expansion to neuronal differentiation and maturation (Figure 2E). Meanwhile, the progressively increased expression of neuronal miRNAs at D120 (including the let-7 family, miR-124-3p, and other neurodevelopmental regulators) further supports the acquisition of mature neuronal identity.

**Figure 2.**
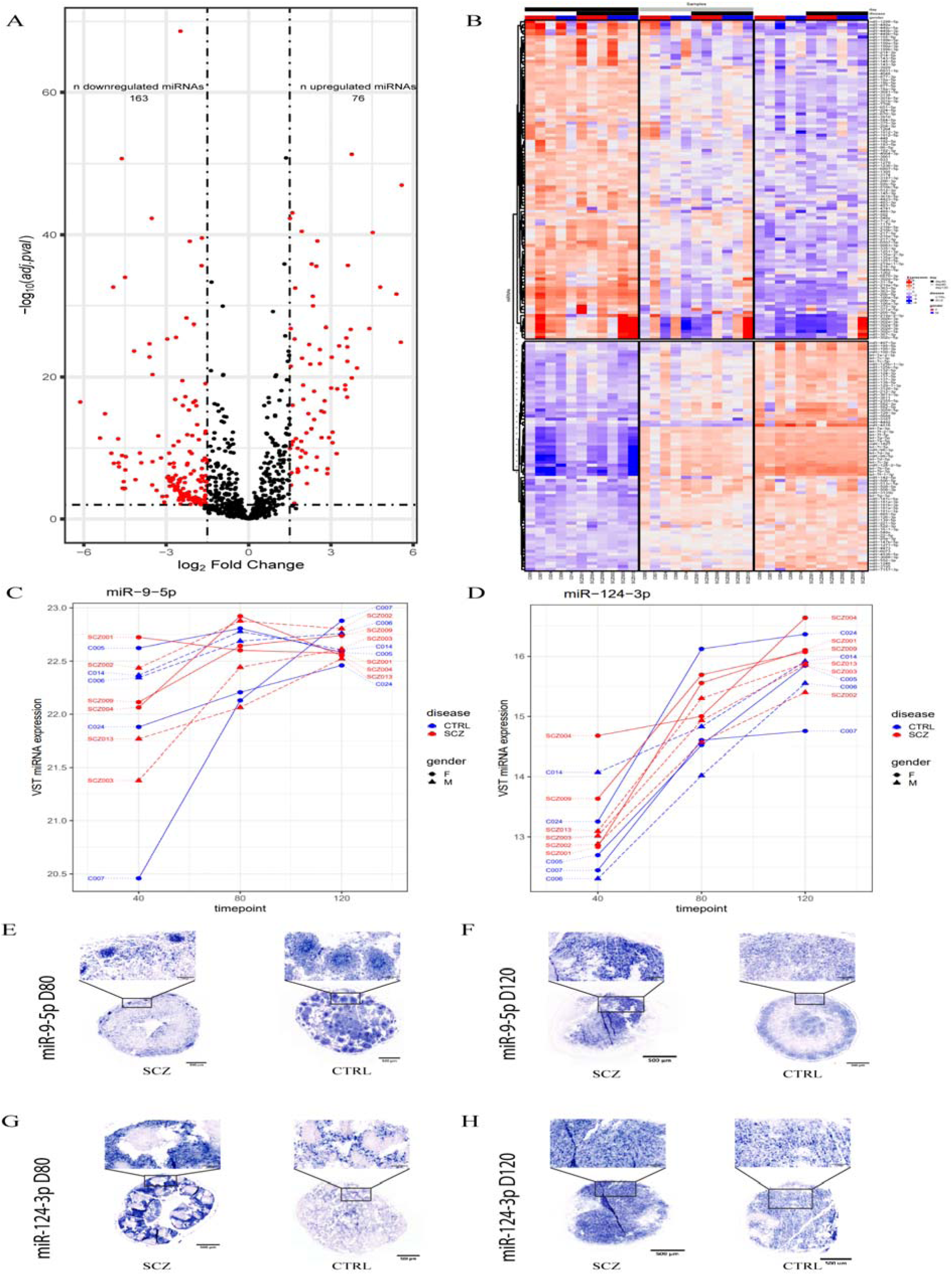
miRNA expression reflects neurodevelopmental progression in SCZ and CTRL DFOs: **A.** Volcano plot showing 163 significantly downregulated and 76 miRNAs (log2FC > ±1.5, p < 0.01) between D40 and D120 DFOs, highlighting dynamic miRNA shifts during neurodevelopment. **B.** Heatmap of differentially expressed miRNAs shows pluripotency/proliferation-associated miRNAs enriched at D40 and neuron-related miRNAs at D120. **C–D**. miR-9-5p(C) and miR-124-3p(D) increase over time, indicating a neurodevelopmental trajectory with hiPSC specific variation. miR-9-5p peaks for most DFOs at D80, indicating a high increase in NPC development. **E–F.** ISH for miR-9-5p at D80 (E) and D120 (F) shows temporal changes in expression during neural differentiation. At D80, miR-9-5p is predominantly localized in NPCs, whereas by D120, expression shifts to mature neurons. This pattern is observed in both SCZ and CTRL DFOs. D80 DFO SCZ001 and C005. D120 DFO SCZ003 and C006. **G–H**. ISH for miR-124-3p shows strong enrichment in neurons at both D80 (G) and D120 (H). This expression pattern is consistent with the established neuronal role of miR-124-3p. D80 DFO SCZ003 and C005. D120 DFO SCZ003 and C005.

Differential expression analysis of miRNAs between D40 and D120 DFOs, irrespective of SCZ or CTRL status, identified 163 significantly downregulated and 76 significantly upregulated miRNAs (Figure 2A; red dots). These reflected DFO maturation with progressive downregulation of miRNAs linked to pluripotency and proliferation and concurrent upregulation of those involved in neural differentiation and development (Figure 2B). Notably, mRNAs encoding proteins involved in synaptic plasticity, neurite outgrowth, cytoskeletal development, and BDNF regulation, which are key processes in neurodevelopment and cognitive function, were predicted targets of several regulatory miRNAs upregulated at D120 [19–21]. Among these, members of the let-7 family showed consistent elevation across all DFOs, in line with their established role in promoting neural differentiation by repressing proliferation-related genes [22]. Similarly, miR-195-5p, miR-497-5p, and miR-16-1-3p were amongst additional miRNAs upregulated at D120 (Figure 2B). Neurodevelopmental miRNAs including miR-128, miR-132, miR-181, miR-212, and miR-582, were elevated at D120, indicating a coordinated shift towards neuronal maturation [23, 24]. Likewise, we observed increased later expression of miR-124-3p, a neuron-specific miRNA essential for proliferation, differentiation, migration, and memory (Figures 2D, G, H). Given that miR-124 accounts for 25-50% of brain miRNAs, its upregulation is consistent with progressive neuronal maturation [25]. miR-9, a neurodevelopmental regulator is highly expressed during this process and balances proliferation and differentiation in embryonic neural progenitors by targeting transcription factors such as TLX and HES1, which maintain progenitor identity, as well as FOXG1, thereby modulating neurogenesis timing and patterning [26, 27]. miR-9 was highly expressed at D40, peaking at D80 (Figure 2C), coinciding with the presence of both progenitors and neurons (Figure 2E). By D120, rosettes disappeared, and mature cell types predominated (Figure 2F), corresponding with declining miR-9 expression as neurogenesis progressed [28].

Thus, DFO maturation is accompanied by dynamic, stage-specific shifts in miRNA expression closely mirroring human neurodevelopment. Initial upregulation of miR-9 and subsequent decline aligns with the transition from progenitor expansion to neuronal differentiation and maturation. Meanwhile, the progressively increased expression of neuronal miRNAs (including the let-7 family, miR-124-3p, and other neurodevelopmental regulators) further supports the acquisition of mature neuronal identity. Our findings confirm that DFOs faithfully recapitulate key molecular features of human cortical development and provide a relevant model for studying the regulatory role of miRNAs in neuronal differentiation and maturation.

### Selective miRNA Dysregulation in SCZ vs. CTRL DFOs

SCZ DFOs exhibit a shift from progenitor proliferation toward premature differentiation, characterized by downregulation of miRNAs that support progenitor maintenance and proliferation and upregulation of miRNAs that promote neuronal differentiation and synaptic maturation. This pattern suggests a truncated developmental window that may underlie early neurodevelopmental abnormalities in SCZ. While SCZ and CTRL DFOs showed broadly similar developmental trajectories (Figures 1–2), differential miRNA analysis revealed 86 upregulated and 173 downregulated miRNAs in SCZ (Figure 3A). The 19 most significantly altered miRNAs (baseMean > 200; adjusted p < 0.1) were validated by qPCR, demonstrating strong concordance with sequencing results (Figure 3B). Nine downregulated miRNAs have established neurodevelopmental functions, including roles in progenitor maintenance, proliferation, and maturation. For example, miR-17-5p and miR-18a-3p, members of the same cluster, are essential for neural progenitor proliferation, and their loss induces premature neuronal maturation [29–31]. In our DFOs, miR-17-5p was enriched in rosettes at D80 (Figures 3C, E) and declined by D120 (Figures 3D, F), consistent with a role in immature neurons. miR-219-2-3p, a brain-enriched regulator of differentiation via repression of negative lineage regulators, was also reduced in SCZ DFOs. TLX sustains NSC proliferation through miR-219 suppression. Accordingly, reduced miR-219 may enhance progenitor proliferation while limiting differentiation, reflecting context-dependent effects that diverge from *in vivo* findings in which elevated miR-219 decreases NPC numbers [32, 33]. In contrast, ten upregulated miRNAs in SCZ are implicated in neuronal differentiation, synaptic development, and apoptosis (Figure 3B). miR-128-3p, miR-101-3p, miR-374a-5p, and miR-361-3p are associated with synaptic plasticity [34–37], whereas miR-30d-5p and miR-187-3p regulate apoptosis and autophagy through BDNF-related pathways, suggesting enhanced cell death and altered autophagy in SCZ DFOs [38, 39]. miR-378a-3p, involved in NSC proliferation via TLX, showed rosette enrichment at D80 (Figures 3G, J) and decreased by D120 (Figures 3H, K), although NGS data indicated a stable increase across time (Figure S6), underscoring the need for multimodal validation.

**Figure 3.**
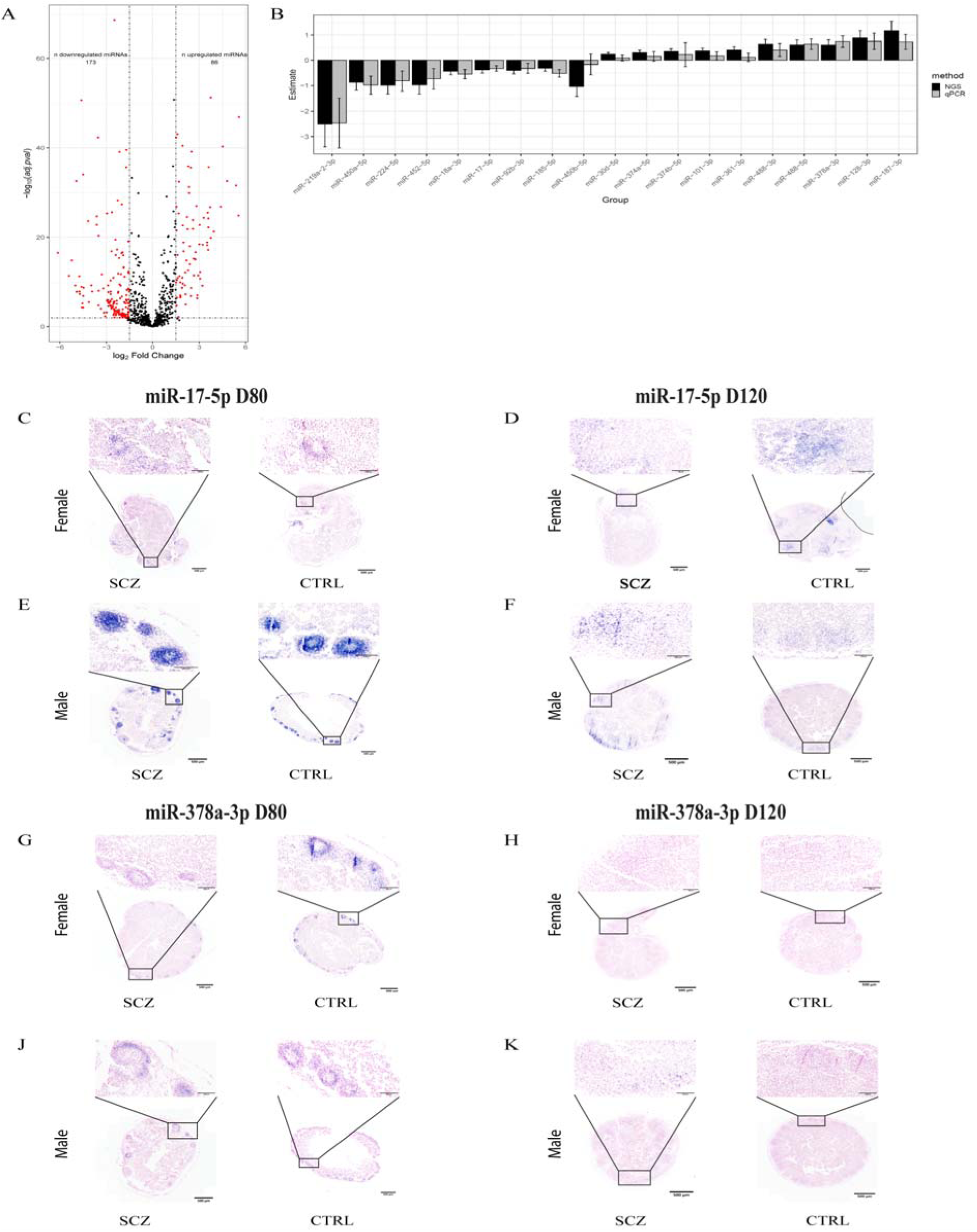
Differentially expressed miRNAs in SCZ vs. CTRL DFOs: **A.** Volcano plot shows 86 upregulated and 173 downregulated miRNAs in SCZ vs. CTRL DFOs (log2FC > ±0.3, adj. p < 0.1). **B.** qPCR validation of 19 miRNAs (basemean >200) confirms expression trends observed in miRNA-sequencing data. **C–F.** ISH staining of miR-17-5p at D80 (C) and D120 (F) reveals nuclear localization in structures resembling neural rosettes, persisting at later stages, with higher expression in male-derived DFOs. Female-derived DFOs show weaker signal. DFOs for Females are SCZ009 and C024 both at D80 and D120. DFOs used for male both at D80 and D120 is SCZ003 and C006. **G–K.** ISH staining of miR-378a-3p shows expression of miR-378a-3p in rosettes at D80 (G), with reduced expression at D120 (K). Expression is stronger in female CTRL DFOs, while male SCZ samples show lower

### Integrated Transcriptomic and Proteomic Analysis Reveals Selective Accelerated Neuronal Differentiation in SCZ DFOs

To assess molecular alterations between SCZ and CTRL DFOs, we integrated transcriptomic and proteomic profiling, focusing on predicted miRNA targets. RNA-seq analysis revealed 52 upregulated and 25 downregulated transcripts in SCZ DFOs (Figure 4A, red dots), including several SCZ risk genes such as *SCG2*, *CACNA1D*, *CAMK2B*, *KCNJ3*, and *EPHA3* [40–44]. Amongst these were *GABRA4*, *CAMK2B*, *EBF1*, and *CDH18*, which were validated by qPCR (Figure 4B). Integrative analysis of matched RNA-seq and proteomics data (6,767 gene–protein pairs) identified 2,628 with strong transcript–protein concordance (Pearson r > 0.5) (Figure 4C). Notably, predicted miRNA targets such as *EPHA3* and *CAMK2B* showed significant and directionally consistent dysregulation (Figure 5D–E). EPHA3 was markedly downregulated at both transcript and protein levels (r = 0.85, adjusted p = 4.5 × 10), consistent with its known role in GABAergic interneuron development through NCAM-mediated RhoA/ROCK signaling [45]. Impaired EPHA3 signaling likely disrupts axon guidance and GABAergic circuit formation, contributing to excitatory–inhibitory (E/I) imbalance in SCZ [45]. In contrast, CAMK2B, encoding the β-subunit of CaMKII, was significantly upregulated (r = 0.91, adjusted p = 1.6 × 10 ¹¹). This finding aligns with elevated CAMK2B expression reported in the SCZ prefrontal cortex [43]. CAMK2B modulates GABA_A receptor function and promotes dendritic maturation and axon guidance [46–48] suggesting that its dysregulation may underlie structural and synaptic abnormalities in SCZ. The strong transcript–protein concordance of dysregulated miRNA targets supports the idea of impaired post-transcriptional regulation by altered miRNAs in SCZ DFOs. Consistent with these molecular changes, SCZ DFOs exhibited features of accelerated, yet dysregulated neurodevelopment compared with controls. GO enrichment analyses of differentially expressed transcripts and proteins revealed upregulation of neuronal signaling and ion channel pathways, alongside downregulation of ribosomal, translational, and chromatin organization processes (Figure 4F–G). Reduced protein expression of pathways related to rRNA metabolism, ribosome biogenesis, and DNA replication further supports premature neurodevelopmental progression [49].

**Figure 4.**
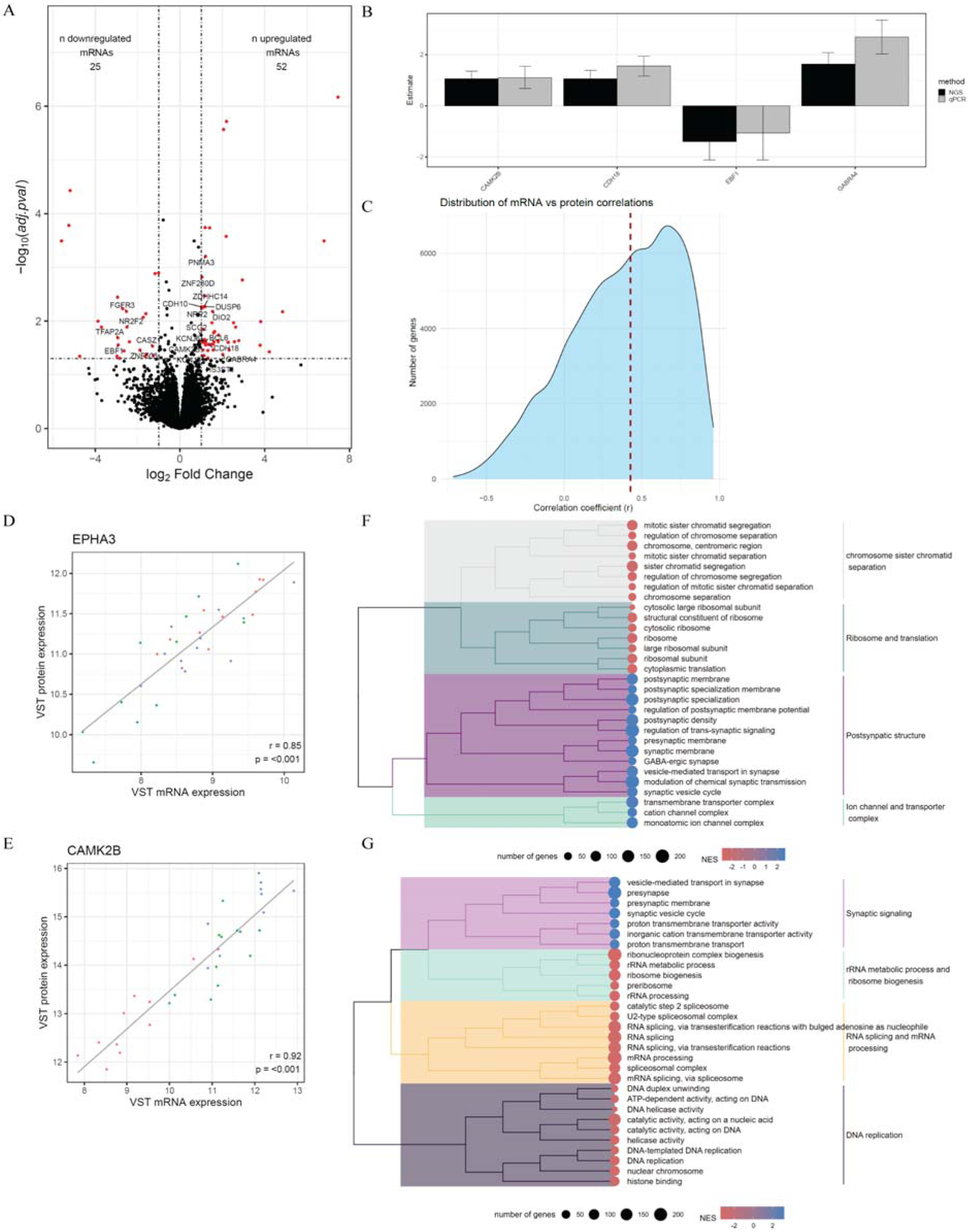
Integrated transcriptomic and proteomic analysis highlights enhanced neurodevelopment in SCZ-derived DFOs: **A.** Volcano plot shows differentially expressed protein-coding genes (mRNAs) (loglFC > ±1, adj. p < 0.05), with highlighted genes targeted by the 19 miRNAs from Figure 4B. **B.** qPCR validation of *GABRA4*, *CAMK2B*, *EBF1*, and *CDH18* confirms trends from mRNA-sequencing data. **C.** Correlation analysis across 6,767 genes shows strong mRNA–protein concordance in 2,628 genes (Pearson r > 0.5). **D.** Examples: EPHA3 (downregulated) and CAMK2B (upregulated) show strong transcript–protein correlation (r = 0.85 and 0.91, respectively). **E-F**. GO enrichment of transcripts and proteins reveals upregulation of neurodevelopmental pathways in SCZ organoids vs. CTRL.

**Figure 5:**
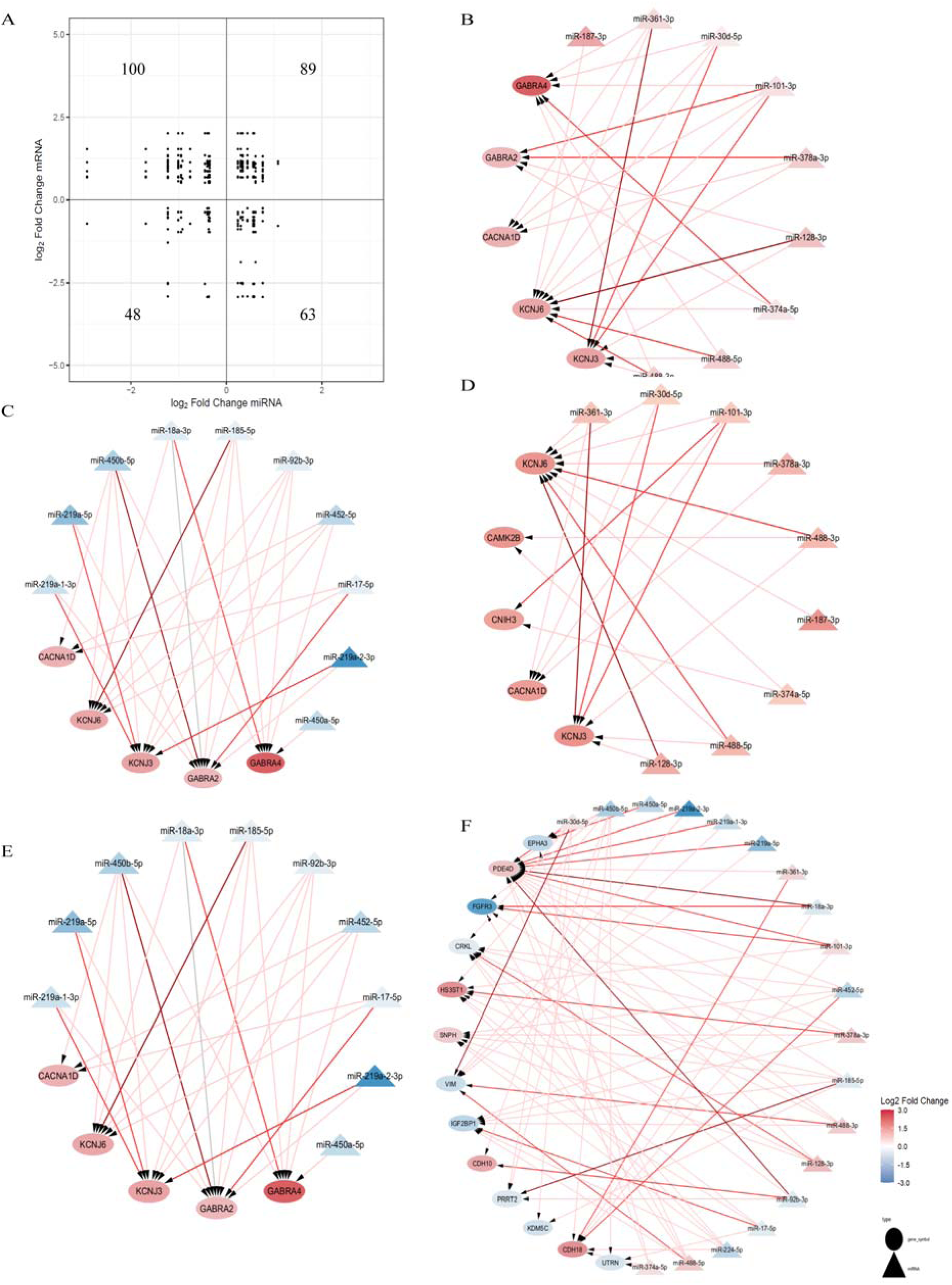
Integrated Analysis of Dysregulated miRNA–mRNA Interactions Across Neuronal Pathways and Synapse Development: **A.** Correlation quadrant plot of significantly dysregulated miRNAs and their predicted mRNA targets. Interactions were predicted using TargetScan (≥1 conserved or non-conserved site) with filters: baseMean >200 for both miRNAs and mRNAs; mRNA padj <0.05, miRNA padj <0.1. The plot shows: 100 interactions were upregulated mRNAs pair with downregulated miRNAs. 63 interactions were downregulated mRNAs pair with upregulated miRNAs. 48 interactions where both mRNAs and miRNAs are downregulated. 89 interactions where both are upregulated. **B.** Upregulated GABAergic pathway genes and their upregulated regulatory miRNAs **C.** Downregulated miRNAs targeting significantly upregulated GABAergic genes **D–E.** Shared upregulated genes across glutamatergic, dopaminergic, and GABAergic neurons with their upregulated (D) and downregulated (E) miRNA regulators **F.** Dysregulated genes involved in synapse development and their interactions with up- or downregulated miRNAs Node colors indicate logl fold change (red = upregulated, blue = downregulated; scale: ±3). Edge colors represent confidence in predicted downregulation: High confidence dark red: score ≤ –0.4, moderate confidence red: –0.4 < score ≤ –0.2, weak confidence light red: –0.2 < score ≤ 0, not regulated: grey

At the protein level, neuronal and synaptic markers including MAP2, MAPT, DCX, and SOX1 were significantly altered (Table S3), and cortical identity markers SATB2 and CTIP2 were upregulated, consistent with immunostaining results indicating accelerated maturation (Figure 1C; Figure S7–S8). GO enrichment additionally highlighted GABAergic pathway dysregulation, with fewer changes in dopaminergic or glutamatergic signaling (Figure 4F; Table S4). This pattern aligns with established models of SCZ pathophysiology implicating E/I imbalance [50–52].

Collectively, the coordinated miRNA, transcriptomic, and proteomic alterations point to premature and dysregulated GABAergic neurodevelopment as a defining molecular signature of SCZ DFOs.

### Dysregulated miRNA-mRNA Interactions Highlight GABAergic and Synaptic Imbalance in SCZ DFOs

To assess how altered miRNA regulation contributes to molecular dysregulation in SCZ DFOs, we examined predicted miRNA–mRNA interactions using TargetScan. Nineteen differentially expressed miRNAs were predicted to target 55 of 77 dysregulated mRNAs, forming approximately 300 potential regulatory pairs (Figure 5A). Among these, 100 interactions involved downregulated miRNAs paired with upregulated mRNAs and 63 involved the reverse pattern, consistent with miRNAs’ repressive role. Instances of concordant regulation (both up- or downregulated) likely reflect indirect or feedback mechanisms.

#### Enrichment of GABAergic and Synaptic Pathways

Functional annotation of target transcripts revealed significant enrichment in GABAergic, glutamatergic/dopaminergic, and synaptic signaling pathways (Figures 5B–F). Notably, *GABRA2* and *GABRA4*, encoding α2 and α4 subunits of the GABA_A receptor, and *KCNJ3*/*KCNJ6*, encoding GIRK1/2 channels essential for inhibitory signaling, were upregulated and predicted targets of multiple downregulated miRNAs. KEGG analysis highlighted increased expression of pre- and postsynaptic GABAergic components, including KCC2 (SLC12A5), a chloride transporter mediating the developmental switch of GABA transmission from excitatory to inhibitory (Figure S9). These results indicate that disrupted miRNA–mRNA regulation contributes to accelerated GABAergic maturation and early (E/I) imbalance, consistent with SCZ pathophysiology [53].

Several miRNAs were predicted to regulate genes involved in neuronal differentiation and synaptic plasticity. miR-101-3p, miR-374-5p, and miR-378a-3p targeted *GABRA2* and *GABRA4*, while miR-128-3p and miR-361-3p targeted *KCNJ3* and *KCNJ6*. Conversely, CACNA1D, encoding a voltage-gated calcium channel critical for synaptic transmission, was upregulated at the transcript but not protein level (Figures 5, Table S3), suggesting post-transcriptional repression by these miRNAs. Downregulation of progenitor-associated miRNAs, including the miR-17-92 cluster, miR-450b-5p, miR-452-5p, miR-185-5p, and miR-219, further supports a shift toward premature neuronal differentiation and an early GABAergic transition, likely contributing to disrupted network maturation in SCZ DFOs.

#### Broader Synaptic and cAMP Pathway Dysregulation

To explore broader functional effects, predicted mRNA targets were mapped to GO and KEGG pathways. Enrichment analyses revealed associations with synaptic membrane components and multiple neurotransmitter systems, including dopaminergic, glutamatergic, serotonergic, and endocannabinoid signaling (Figures 6A–B). Among genes involved in synaptic development, *CDH10*, *CDH18*, and *PDE4D* involved in cell adhesion, axon guidance, and neuronal plasticity, were strongly regulated (Figure 5F). *PDE4D*, targeted by 16 distinct miRNAs, emerged as a major regulatory hub controlling the balance between progenitor proliferation and neuronal differentiation. Both *PDE4D* and its paralog *PDE4B*, previously implicated in major mental illness [54, 55], belong to the phosphodiesterase-4 (PDE4) family, which modulates cAMP signaling, a pathway critical for neuronal development and cognitive function [56].

**Figure 6:**
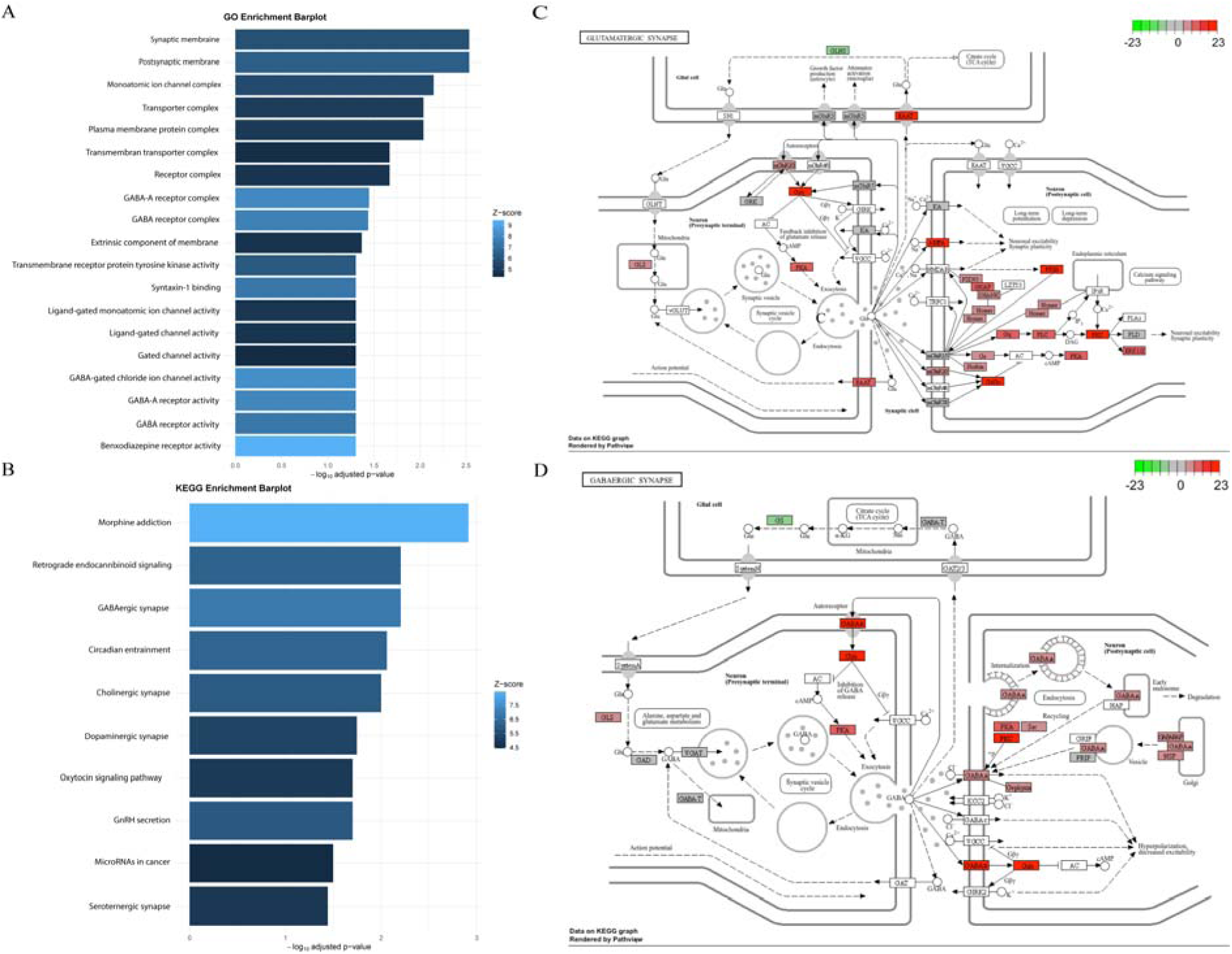
miRNA–mRNA–Pathway interaction network and synaptic protein alterations in SCZ organoids: **A.** Enrichment GO terms on the genes (mRNAs) that are targeted by differentially regulated miRNAs in SCZ vs. CTRL, (adj. p < 0.05) **B.** Subset showing only KEGG-mapped genes targeted dysregulated miRNAs, highlighting top SCZ-relevant pathways and potential miRNA influence during early brain development, (adj. p < 0.05) **C.** Altered expression of glutamatergic synapse proteins and receptors in SCZ organoids; red = upregulated, green = downregulated. **D.** Dysregulated proteins in GABAergic synapses show altered expression of GABAB receptor and other key synaptic components in SCZ; red = upregulated, green = downregulated

#### Excitatory Synaptic Compensation

Although transcript-level changes in excitatory synapse genes were limited, several glutamatergic proteins were significantly upregulated, including GRIA2, SHANK, and PSD95 (Figure 6C; Table S3). These increases suggest reduced miRNA-mediated repression and compensatory upregulation of excitatory synaptic proteins in response to GABAergic dysfunction. The involvement of SHANK and PSD95, previously linked to SCZ susceptibility [57], indicates coordinated dysregulation of both inhibitory and excitatory synapses, consistent with an altered developmental trajectory of cortical network assembly.

Together, these results demonstrate that dysregulated miRNA–mRNA networks in SCZ DFOs converge on pathways governing GABAergic maturation, synaptic organization, and neuronal differentiation. Reduced miRNA mediated repression of key inhibitory and ion channel genes (e.g., *GABRA2, GABRA4, KCNJ3, KCNJ6*) likely drives premature GABAergic development and E/I imbalance, while compensatory upregulation of excitatory synaptic components further exacerbates network dysregulation (Figure 5A,B and 6D). These findings identify altered post-transcriptional regulation as a mechanistic link between miRNA dysfunction and the neurodevelopmental abnormalities underlying SCZ.

#### Dysregulation of Cell Adhesion, Cytoskeletal, and ECM Pathways Suggests Structural Connectivity Deficits in SCZ DFOs

In addition to its established role in synapse development, *HS3ST1*, which encodes heparan sulfate glucosamine 3-O-sulfotransferase 1, is critical for neuronal circuit formation through interactions with cell-surface and extracellular matrix (ECM) proteins and related signaling pathways [58, 59]. In SCZ DFOs, *CDH10* and *CDH18*, members of the cadherin family mediating calcium-dependent cell–cell adhesion and interacting with integrins at the ECM interface, were upregulated (Figure 5F). This may represent a compensatory response aimed at stabilizing synaptic contacts and maintaining connectivity during neurodevelopment [60].

Conversely, several genes essential for cytoskeletal organization and ECM remodeling, including VIM (Vimentin), UTRN (Utrophin), and CRKL (Figure 5F), were downregulated. These proteins support cell migration, cytoskeletal anchoring, and ECM maintenance [61–63]. Their reduced expression may compromise cellular motility and structural stability, potentially contributing to ECM abnormalities and disrupted cortical organization observed in SCZ [64].

Integrative miRNA–mRNA network analysis revealed that *HS3ST1*, *CDH10*, and *CDH18* were predicted targets of downregulated miRNAs, including the miR-17-92 cluster, miR-224-5p, and miR-450b-5p, whereas *VIM*, *UTRN*, and *CRKL* were predominantly targeted by upregulated miRNAs such as miR-30d-5p, miR-128-3p, miR-488-3p, and miR-488-5p (Figure 5F). This inverse regulation supports a model in which miRNA dysregulation disrupts the coordinated control of cell adhesion, cytoskeletal organization, and ECM remodeling, thereby impairing neuronal migration and structural connectivity during cortical development in SCZ.

## DISCUSSION

Our findings support the hypothesis that dysregulated miRNA networks contribute to SCZ-associated neurodevelopmental abnormalities. SCZ-derived DFOs displayed accelerated neuronal maturation and broad transcriptomic and proteomic changes consistent with aberrant regulation of neuronal differentiation, synaptic signaling, and E/I balance [50, 65, 66]. These alterations likely reflect, at least in part, the impact of disrupted miRNA expression on key developmental signaling pathways.

### miRNA Dysregulation and Neurodevelopmental Timing

SCZ DFOs showed downregulation of miRNAs involved in neural progenitor proliferation alongside upregulation of neurodevelopment-associated miRNAs. Reduced miR-92a/b expression, linked to PI3K/AKT, TGF-β, and WNT signaling [67, 68], suggests impaired progenitor maintenance and disrupted coordination of neurodevelopmental cascades [69]. Downregulation of miR-92b-3p likely reflects premature or accelerated neuronal differentiation, consistent with structural and maturation abnormalities reported in SCZ. Conversely, miR-9 and miR-124-3p were progressively upregulated in both groups, reflecting their established roles in promoting neuronal differentiation and maturation [26, 70, 71]. Together, these findings validate the temporal fidelity of our DFO model in recapitulating key developmental transitions.

Interestingly, miR-137, a well-established SCZ risk-associated miRNA, did not differ between SCZ and control DFOs. Given prior reports that miR-137 overexpression alters synaptic ultrastructure rather than specific protein abundance [72], its lack of differential expression here may reflect donor genotype, human-specific regulation, or later developmental effects beyond D120.

### Altered miRNA–mRNA Networks Underlie GABAergic and Synaptic Imbalance

Integration of miRNA and mRNA data revealed 300 predicted regulatory interactions, 163 of which displayed canonical inverse relationships, highlighting widespread post-transcriptional dysregulation. These networks converged strongly on GABAergic signaling pathways, implicating premature inhibitory circuit maturation in SCZ DFOs. Multiple downregulated miRNAs: miR-101-3p, miR-378a-3p, miR-374-5p, miR-361-3p, and miR-128-3p, targeted GABA receptor subunits (*GABRA2*, *GABRA4*), GIRK channels (*KCNJ3*, *KCNJ6*), and the chloride transporter KCC2, which mediates the developmental switch from depolarizing to hyperpolarizing GABA [73]. Upregulation of these genes suggests premature GABAergic maturation, consistent with early E/I imbalance implicated in SCZ [50].

Beyond inhibitory signaling, gene ontology and pathway analyses revealed broader alterations across glutamatergic, dopaminergic, serotonergic, and endocannabinoid pathways, consistent with system-wide synaptic dysregulation [74–77]. Notably, *PDE4D* emerged as a central regulatory hub, targeted by 16 dysregulated miRNAs. Given its role in neuronal differentiation, cognitive function and its genetic association with major mental illness [78], *PDE4D* dysregulation likely contributes to disrupted progenitor maintenance and premature neuronal maturation via altered cAMP signaling [79].

At the protein level, SCZ DFOs exhibited increased abundance of AMPA receptor subunits (GRIA2) and postsynaptic scaffolding proteins SHANK and PSD95 [57, 80, 81] suggesting reduced miRNA repression may enhance excitatory synaptic protein synthesis. This could represent a compensatory mechanism for GABAergic dysfunction. Upregulation of *CAMK2B* and *CNIH3*, both key regulators of excitatory synapse formation, further underscores excitatory circuit remodeling in SCZ DFOs.

### Structural and Extracellular Matrix Dysregulation

Beyond synaptic networks, we identified consistent dysregulation of genes involved in cell adhesion, cytoskeletal organization, axon guidance, and ECM remodeling pathways increasingly recognized as contributors to SCZ [60, 82, 83]. Downregulation of VIM, CDH10, and CDH18, together with decreased expression, points to compromised structural integrity, impaired neuronal migration, and disrupted connectivity during cortical development. In contrast, upregulation of EPHA3, a receptor tyrosine kinase involved in axon guidance, may reflect compensatory remodeling or aberrant circuit wiring.

The ECM provides the structural framework for neuronal migration, synapse formation, and circuit stabilization; thus, its dysregulation could contribute to the early neurodevelopmental disturbances observed in SCZ. Given its accessibility and dynamic nature, the ECM represents a promising therapeutic target. Inhibitors of ECM-degrading enzymes such as MMP-9, which have shown benefit in amyotrophic lateral sclerosis and Alzheimer’s disease [84, 85] could potentially restore neuronal architecture and connectivity in SCZ, particularly in treatment-resistant or developmentally driven subtypes.

In summary, our findings demonstrate that dysregulated miRNA–mRNA networks in SCZ DFOs converge on pathways controlling neuronal maturation, synaptic organization, and ECM integrity. These alterations collectively suggest premature neuronal differentiation, disrupted E/I balance and impaired structural connectivity as early mechanistic features of SCZ pathophysiology. By integrating miRNA, transcriptomic, and proteomic data within a human organoid model, this work highlights miRNA-mediated post-transcriptional regulation as a unifying mechanism linking molecular, cellular, and circuit-level disturbances in SCZ.

## STRENGTH and LIMITATIONS

Our study has several notable strengths. By using hiPSC-derived DFOs, we leveraged a human-specific model that recapitulates essential molecular and spatial features of cortical development, enabling direct investigation of early neurodevelopmental processes that cannot be accessed through postmortem or *in vivo* approaches. The integration of miRNA profiling with transcriptomic and proteomic analyses provided a comprehensive multi-omics framework, allowing us to identify convergent regulatory networks and strengthening the inference of miRNA–mRNA interactions implicated in SCZ. Immunohistochemistry, and in situ hybridization further validated the cellular context of dysregulated molecules, adding spatial resolution to the molecular findings. Importantly, the use of multiple independent SCZ and control hiPSC lines supported the robustness and reproducibility of the observed signatures.

Several limitations should also be considered. Organoids inherently exhibit line-to-line and batch variability, which, although mitigated by standardized differentiation protocols, may introduce biological noise. DFOs also lack certain cell populations found in the developing cortex, including microglia and vascular elements, potentially restricting the range of disease-relevant interactions that can be modeled. Additionally, organoids primarily capture early to mid-gestational developmental stages, limiting insights into later phases of cortical maturation where additional SCZ-related alterations may arise. miRNA–mRNA relationships identified here are based on predictive and correlative analyses, and functional perturbation experiments will be necessary to establish causal regulatory mechanisms. Finally, the relatively modest sample size, typical for organoid-based studies, may limit the detection of more subtle or heterogeneous disease-associated effects.

## CONCLUSIONS and PERSPECTIVES

Our findings demonstrate that dysregulated miRNA–mRNA networks in SCZ DFOs converge on pathways controlling neuronal maturation, synaptic organization, and ECM integrity. These alterations collectively suggest premature neuronal differentiation, disrupted E/I balance and impaired structural connectivity as early mechanistic features of SCZ pathophysiology. By integrating miRNA, transcriptomic, and proteomic data within a human organoid model, miRNA-mediated post-transcriptional regulation emerges as a unifying mechanism linking molecular, cellular, and circuit-level disturbances in SCZ.

## Author contributions

FAM and KF conceived the idea for this study. FAM designed the experiments and interpreted the data. RS supported by FAM analyzed the miRNA and mRNA sequencing data. PJ, AN and MRL performed proteomics. RS and FAM incorporated proteomics data into miRNA and mRNA results. SC advised on qPCR for miRNA validation. CBA performed CNV analyses and interpretation of the hiPSC. MB was responsible for and supervised the patient recruitment. EB recruited, assessed patients and retrieved blood samples. BSN advised on miRNA ISH. MW, TM and MK assisted with experimental procedures. FAM and KF wrote the manuscript with input from all the other authors. All authors have approved the final version. All data needed to evaluate the conclusions in this article are present in the article and/or the supplemental materials. The authors report no biomedical financial interests or potential conflicts of interest.

## Data sharing

Due to potential personal identifiable information, data are shared on a group level only with summary statistics.

